# Interferon-alpha2 but not Interferon-gamma serum levels are associated with intramuscular fat in obese patients with nonalcoholic fatty liver disease

**DOI:** 10.1101/384297

**Authors:** Giovanni Tarantino, Susan Costantini, Vincenzo Citro, Paolo Conforti, Francesca Capone, Angela Sorice, Domenico Capone

**Affiliations:** Department of Clinical Medicine and Surgery, “Federico II” University Medical School of Naples, Italy; Oncology Research Center of Mercogliano (CROM), Istituto Nazionale Tumori – IRCSS – Fondazione G. Pascale, Napoli, Italy; Department of General Medicine, “Umberto I” Hospital of Nocera Inferiore (ASL Salerno), Italy; “Federico II” University Medical School of Naples, Italy; Integrated Care Department of Public Health and Drug-Use, Section of Medical Pharmacology and Toxicology, “Federico II” University, Naples, Italy.

**Keywords:** IFN-alpha, IFN-gamma, MTG, obesity, NAFLD

## Abstract

**Background:** Intramuscular triglycerides (IMTGs) represent an important energy supply and a dynamic fat-storage depot that can expand during periods of elevated lipid availability and a fatty acid source. Ultrasonography (US) of human skeletal muscles is a practical and reproducible method to assess both IMTG presence and entity.

Although a crosstalk between cytokines in skeletal muscle and adipose tissue has been suggested in obesity, condition leading to hepatic steatosis (HS) or better defined as nonalcoholic fatty liver disease and cancer, there are still questions to be answered about the role of interferons (IFNs), alpha as well as gamma, and IMTG in obesity. We aimed at discovering any correlation between IFNs and IMTG.

**Methods:** We analysed anthropometric data, metabolic parameters and imaging features of a population of obese subjects with low-prevalence of co-morbidities but HS. The levels of serum IFNs were detected by a magnetic bead-based multiplex immunoassays.

**Results:** Serum concentrations of IFN-alpha2 were increased, while serum levels of IFN-gamma were decreased confronted with those of controls; the severity of IMTG, revealed at US as Heckmatt scores, was inversely predicted by IFN-alpha2 serum concentrations; IMTG scores were not predicted by serum levels of IFN-gamma; IMTG scores were predicted by HS severity, ascertained at US; HS severity was predicted by visceral adipose tissue, assessed by US, but the latter was not instrumental to IMTG.

**Discussion & Conclusion:** This study has added some pieces of observation about the cytokine network regulating the interplay between IMTG and obesity in obese patients with HS.

## INTRODUCTION

Intramuscular fat, also known as intramuscular triglycerides, intramuscular triacylglycerol or intramyocellular triacylglycerol (IMTG), but also intramuscular adipose tissue (IntraMAT) and intramyocellular lipid (IMCL), when increased is thought to be linked to increased lipolytic activity in skeletal muscle, contributing to inducing insulin resistance (IR), (1). Increased muscle TG stores are characterised by cytosolic accumulation of diacylglycerol and acyl-CoA-triglycerides, being lipase regulation central to skeletal muscle lipolysis (2). Disturbances in pathways of lipolysis may play a role in the development and maintenance of these increased fat stores. Not only is IMTG an important energy supply for skeletal muscle, but represents a dynamic fat-storage depot that can expand during periods of elevated lipid availability (3). Structural characteristics of IMCL seem to be similar between highly trained endurance athletes, type 2 diabetes patients, and overweight, sedentary men after an overnight fast. This observation is not in agreement with the hypothesis that elevated IMCL deposits are direct responsible for inducing IR (4).

A recent study, carried out to evaluate the exact localisation of IntraMAT, using ^1^H magnetic resonance spectroscopy or echo intensity (EI) determined by B-mode ultrasonography (US) of human skeletal muscles, has surprisingly suggested that IntraMAT primarily reflects extra-myocellular lipids, not IMCL (5).

Focusing on techniques unravelling IMTG, i.e., EI at US and high-resolution T1-weighted MRI, strong correlations were found between MRI percent fat and muscle EI after correcting for subcutaneous fat thickness (6).

But, apart evidence for being muscle US a practical and reproducible method, another point to be cleared consists in the choice of muscular district to be explored, i.e., the location of the region of interest. Here again, research has confirmed that the EI of biceps brachii and tibialis anterior was higher than that of all other muscles (7).

Beyond considerable insight into the role of IMTG in acute and chronic exercise training (8) and according to the proposed crosstalk between myokines/cytokines and adipokines in skeletal muscle and adipose tissue (9), there are still questions to be answered about the link of interferons (IFNs), alpha as well as gamma, and IMTG in obesity. Indeed, some data show that IFN-gamma, which is released from inflamed omental adipose tissue, may contribute to the metabolic abnormalities seen in human obesity (10). What is more, investigation pointed out to increased levels of IFN-gamma in obese subjects that were associated with central adiposity (11).

Indeed, little research has been conducted to date on the role of IFN-alpha on visceral fat excess (12) and none on IMTG. The subtype 2 of IFN-alpha was chosen to be evaluated in this study due to its action on memory CD8 cells and cytotoxic CD8 cells, which are activated by adipose tissue, in turn promoting the recruitment and activation of macrophages in this tissue (13). Still, according to literature data, obese subjects showed a decreased ability to produce IFN-alpha in response to Toll-like-receptors ligands, partially explaining their decreased response to viral infections (14) Aiming at finding any correlations between serum concentrations of IFN-alpha2 as well as IFN-gamma and IMTG, we analysed a population of obese subjects with low-prevalence of co-morbidities but nonalcoholic fatty liver disease (NAFLD) or hepatic steatosis (HS), evaluated by US.

Finally, there is no fresh evidence corroborating the link between NAFLD and IMTG outside exercise intervention (15), taking into account that IMTG is linked to increased BMI and visceral obesity (16) and shares common mechanisms with NAFLD.

## METHODS

### Patients

We carried out a cross sectional type of observational study where at a particular point of time we described characteristics of obese patients without follow-up, with main variables, i.e., IFNs levels and IMTG scores compared to controls.

Specifically, this sub-study used the same original patient sample contained in a previous research (17), but with completely different analytical approaches resulting to be equally valid, according to The International Committee of Medical Journal Editors (ICMJE) at http://www.icmje.org/recommendations/browse/publishing-and-editorial-issues/overlapping-publications.html Were selected 80 patients, which fulfilled the inclusion criteria and had given previous oral or, when possible, written consensus, plus 38 healthy subjects, as control group.

### Inclusion criteria

Obese patients of different grade of obesity, on calorie-reduced, low-fat diet, with low prevalence of co-morbidities, such as type 2 diabetes mellitus and hypertension but NALFD, US-documented.

### Exclusion criteria

Patients were excluded if, at the time of blood specimen collection, they self-reported present or antecedent (past month) influenza or cold status, or there had been a history of unexplained weight loss in the past months (i.e., ± 10% initial body weight) or recent illness/chronic disease, and the use of supplements or medications that might have affected body composition or muscle metabolism (eg, steroids). Sarcopenic obese were ruled out from this selection.

Furthermore, any viral, autoimmune, metabolic liver disease (Wilson disease, hemochromatosis or antitrypsin deficiency) was ruled out by using appropriate testing, according to well-accepted diagnostic guidelines. Celiac disease was excluded by evaluating IgA anti-tissue transglutaminase antibodies. Alcohol abuse was disallowed, following the DSM-IV diagnostic criteria, by means of screening tests such as MAST (Michigan Alcohol Screening Test) and CAGE (Cut down, Annoyed, Guilty, and Eye opener), as well as random tests for blood alcohol concentration and the use of a surrogate marker, e.g., Mean Corpuscular Volume. Patients on antihypertensive drugs, and those treated with metformin or insulin, maintained a balanced therapeutic regimen throughout the study.

### Anthropometric evaluation

The three degrees of obesity (light, moderate, and severe or 1-2-3) were established on the basis of BMI cut-off points of 30–34.9 and 35–39.9 and >40□kg/m2, respectively. Visceral obesity was identified by measuring WC at the midpoint between the lower border of the rib cage and the iliac crest. Hip circumference was measured around the widest part of the buttocks, with the tape parallel to the floor, and the waist-to-hip (W/H) ratio was calculated.

### Metabolic profile

The canonical Adults Treatment Panel III was chosen to define the metabolic syndrome, considering at least three criteria: plasma glucose concentrations ≥100 mg dL-^1^, WC >102/88 cm (male/female), serum HDL concentration <50 mg dL^−1^ for women and <40 mg dL^−1^ for men, blood pressure ≥130/85 mm Hg, and serum triglyceride concentration ≥150 mg dL^−1^. Triglyceride values of subjects who had fasted at least 12/14 h before the blood draw were evaluated, averaging the results of at least two determinations, made on different days.

### Laboratory assessment

IFNs levels of 78 patients derived by a previously studied 48-cytokine/chemokine panel (17), which was performed on serum samples using a magnetic bead-based multiplex immunoassays (Bio-Plex) (BIO-RAD Laboratories, Milano, Italy) following manufactures’ instructions. Data from the reactions were acquired using the Bio-Plex 200 reader, while a digital processor managed data output and the Bio-Plex Manager software returned data as Median Fluorescence Intensity (MFI) and concentration (pg/mL). Insulin resistance was studied by the HOmeostatic Metabolic Assessment (HOMA) method with the formula: fasting insulin (μU/mL) x fasting glucose (mg/dL)/405 (18). More than five determinations of HOMA in different situations were taken into account. HOMA-derived β-cell function (HOMA-%B) was also calculated, using the following formula: 20 × fasting insulin (μU/ml)/fasting glucose (mmol/L) − 3.5 (18). A stringent value of HOMA >2 was introduced as limit of the presence of insulin resistance (19). We calculated a quantitative insulin sensitivity check index (QUICKI) as 1/[log(fasting insulin μU/ml + log(fasting glucose mg/dL]), with range between 0.45 in healthy individuals and 0.30 in diabetics (20).

### Ultrasonography features

US measurements were obtained by an Esaote (Genoa, Italy) system. The classification of “bright liver” or hepatic steatosis (HS) was based on the following scale of hyperechogenity: 0 = absent, I = light, 2 = moderate, 3 = severe, pointing out the difference between the densities of the liver and the right kidney (21), using a Convex Probe, with access to the liver through intercostal spaces along the mid-axillary line.Transverse scanning was performed to measure the subcutaneous adipose tissue (SAT) and visceral adipose tissue (VAT) using an eleven and 3.5□ MHz linear probe convex probe, respectively. The SAT was defined as the thickness between the skin-fat interface and the linea alba, avoiding compression, evaluated at the superior tertile of xifo-umbelical line. The VAT was defined as the distance between the anterior wall of the aorta and the internal face of the rectoabdominal muscle perpendicular to the aorta, measured one cm above the umbilicus. When the aortic walls were not visualized as they were obscured by bowel gas, the Doppler scan was used (22).

Muscle US, performed at the level of the biceps brachii of the left superior arm, is a convenient technique to visualise pathological muscle tissue, as it provides results in real time. Both infiltration of fat and fibrous tissue increase muscle echo intensity; that is, the muscles become whiter on the ultrasound image (23). To describe muscle echo intensity, Heckmatt and coworkers developed a visual grading scale in which grade I represented normal muscle and grade IV represented a severely increased muscle echo intensity with total loss of bone echo (we chose biceps brachii versus humerus (24). The levels of brightness of the liver and the biceps brachii, obtained by a single traverse image, were calculated three times directly from the frozen images. The choice of evaluating single traverse image findings was made according to Jenkins et al, who demonstrated that a single transverse imaging and panoramic US imaging are comparable (25).

### Indirect Calorimetry

RMR was measured by indirect calorimetry using a canopy system (V max 29 N, Sensor Medics, Anaheim, USA) in a quiet environment and with patients in the supine position for 30□min before measurement. After a 15–20□min adaptation to the instrument, oxygen consumption and carbon dioxide production were determined for 45□min. Energy expenditure was derived from CO2 production and O2 consumption with the appropriate Weir formula neglecting protein oxidation. (26) BMR, expressed as kcal/24□h, was adjusted for changes in fat-free mass (FFM), which was evaluated by single-frequency bioimpedance analysis (BIA) obtaining a RMR/FFM ratio, expressed as kcal/24□h*kg of body. Fat mass and FFM % were estimated using the device’s standard built in prediction equations and were displayed on the machine and printed out (27). The BIA assessment was performed between 10:00 AM and 4:00 PM. The participants were required to fast and avoid vigorous exercise for at least 1h before BIA assessment. The measurements were recorded by well-trained staff.

Sarcopenic obesity was defined minus of two lower quintiles of muscle mass(<9.12 kg/m^2^ in men and <6.53 kg/m^2^ in women) and two highest quintiles of fat mass (>37.16% in men and > 40.01% in women according to NHANES II (28).

### Control group

Though IFN-alpha2 is one of cytokines/chemokines not detected in any group of age of healthy subjects, either because they are under the lower limit of detection or because they are not produced (29) we took into account values of a population of 38 young subjects to reduce type I error. The control arm provided information when analysing difference of IFN-gamma levels in groups, too.

### Statistics

Data, derived from a normally distributed population were given as mean plus SD. Variables not normally distributed or ordinals are expressed as median (25-75 IQR). The difference in medians was assessed by the Mann-Whitney test. The Two-Way crosstabulation was set by the Pearson correlation coefficient (chi square). Spearman’s coefficient of rank correlation (rho) was employed to analyse the basic correlation between some data.

At univariate analysis, the linear regression analysis (ordinary least squares or OLS) was used evaluating the coefficient with its standard error, 95% confidence intervals (CI) and the t (t-value) and R^2^. In suspicion of heteroscedasticity, i.e., when there were sub-populations that have different variabilities from others in the homoscedastic model, and having detected the presence of few outliers, we analysed the correlation by the robust regression, using Least Absolute Deviations (LAD) Regression. Contextually was conducted a residual analysis, a “residuals versus fits plot”. It is a scatter plot of residuals on the y axis and fitted values (estimated responses) on the x axis. The plot was used to detect non-linearity, unequal error variances, and outliers.

A simultaneous quantile regression was applied as a way to discover more useful predictive relationships between variables (bootstrap method). Quantile regression is more robust to non-normal errors and outliers.

At multiple linear regression also the factor Beta (β) was added.

An ordered probit model was employed to estimate relationships between an ordinal dependent variable (IMTG) or HS at US and a set of independent variables.These ordinal variable are variables that are categorical and ordered, expressed as severity score (I-IV) for the former or severity grade for the latter (1-3). The output showed the coefficients, their standard errors, the z-statistic (also called a Wald z-statistic), and the associated p-values. In a specific circumstance a Bayesian inference computed the posterior probability, expressed as mean, SD, Montecarlo standard error or MCSE, median and credibility intervals.

To highlight light unobserved confounding variables two methods were adopted: j)Testing for mediation was performed as a four step approach in which several regression analyses were performed; the significance of the coefficients were examined at each step to study the so-called indirect effect (30). jj)The method of **I**nstrumental Variables (IV) was utilised to estimate causal relationships. A valid instrument induces changes in the explanatory variable (covariate) but has no independent effect on the dependent variable, allowing to uncover the causal effect of the explanatory variable on the dependent variable. An instrument is a variable that does not itself belong in the explanatory equation but is correlated with the endogenous explanatory variables, conditional on the value of other covariates. The type of model was random effects and the estimator was the Baltagi-Changone.

The Factor Analysis was applied to detect the structure in the relationships among variables, selecting a subset of variables having the highest correlations with the principal component factors. In order to select a subset of variables, firstly Cattell Screen plot, with relative eigenvalues, was performed to screen the real factors, which resulted to be three. Secondly, extraction of the main components amounted to a variance maximizing (varimax) the rotation of the original variable space. The critical value was calculated by doubling Pearson’s correlation coefficient for 1% level of significance (5.152)/square root of patients minus 2 (n 78), i.e., 0.583. In bold will be shown the main components for any single factor, with a value superior to the critical one.

A closed form estimator of the uniqueness (unique variance) is proposed. It has analytically desirable properties, i.e., consistency, asymptotic normality and scale invariance. The concordance correlation coefficient (ρ_c_), which measures precision and accuracy, was adopted to evaluate the degree of pair observations at US.

Stata 15.1, Copyright 1985-2017, was the program on which we run statistics.

## RESULTS

### Characteristics of the selected population are shown in Table 1

The median plus IQR for IFN-alpha2 of healthy subjects was 2 pg/ml (0-2), the age-related reference intervals is shown in Figure 1 of supplemental data.

**Figure 1:**
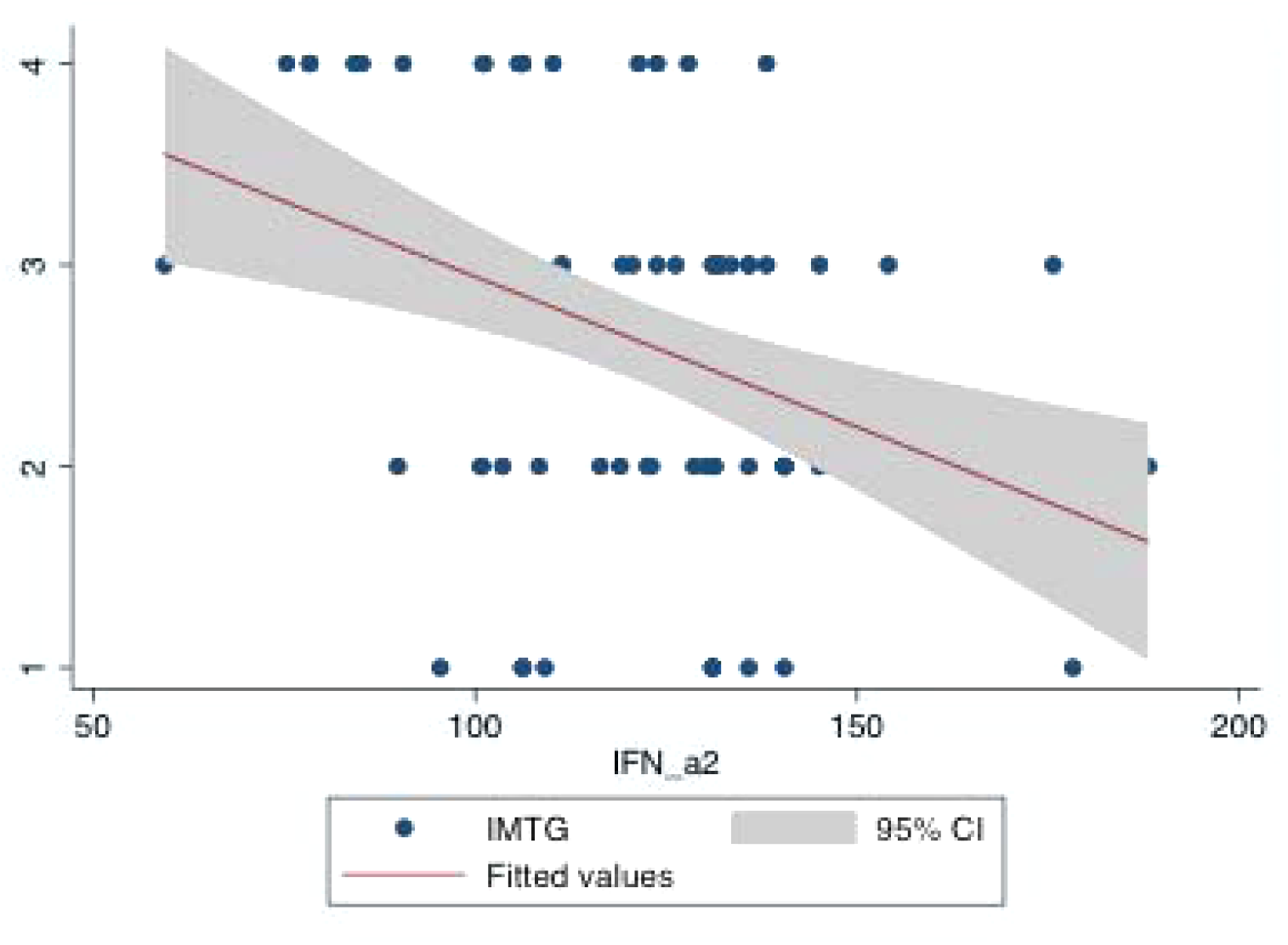
Prediction of IMTG scores by IFN-alpha concentrations. Legend to Figure 1: R^2^ = 0. 1486. This low R-squared graph shows that even noisy, high-variabcan have a significant trend. The trend indicates that the predictor variable still provides information about the response even though data points fall further from the regression line

An interesting finding was that IFN-alpha2 levels of obese patients were found to be significantly increased when compared to those of controls, i.e., 121.9 pg/mL (103.5-135.8) versus 2 (0-2), median plus IQR, P=>0.0001, the Mann-Whitney test. The median plus IQR for IFN-gamma of healthy subjects was 498 pg/ml (468-640), the age-related reference intervals is shown in Figure 2 of supplemental data.

**Figure 2: Prediction of IMTG scores by HS at US grades Legend to Figure 2: R^2^ = 0. 1028; Also this low R-squared graph shows that even noisy, high-variability data can have a significant trend. The trend indicates that the predictor variable still provides information about the response even though data points fall further from the regression line**.

The obesity degrees (1, 2 and 3), to which belonged 7, 26 and 45 patients respectively, did non show different distribution of IFN-alpha levels, Pearson’s chi square, P = 0. 66. Noteworthy, IFN-gamma levels in obese were lower than those of healthy subjects, i.e., 158 pg/mL (56-390) versus 547.5 pg/mL (479-570), median plus IQR, p<0.001, the Mann-Whitney test.

IMTG presence in our population was characterised by a light-moderate score of severity, as reported in Table 1.

**Table 1:** Data of studied patients. Legend to Table 1 # 78 patients were examined for IFNs, Intramuscular TriGlycerides, IMTG; Visceral Adipose Tissue,VAT; Subcutaneous Adipose Tissue, SAT; UltraSound, US; Waist-To-Hip Ratio, WHR; Waist Circumference, WC; Resting Metabolic Rate, RMR; Fat-Free Mass, FFM; Metabolic Syndrome, MS; Hepatic Steatosis, HS;

|  |  |  |  |
| --- | --- | --- | --- |
| Age (yrs) | 46(34-53) | Gender M/F (n) | 36/44 |
| BMI | 42(38-47) | WC cm | 121(114-133) |
| WHR | 0.96(0.94-0.99) | Obesity<br>grade I/II/III (n) | 8/26/46# |
| HOMA | 2.78(1.85-4.18) | HOMA%B | 37.11(22.449.8) |
| QUICKI | 0.32<br>(0.31-0.35) | Insulin<br>μU/mL | 11(7.1-15.8) |
| HDL males<br>mg/dL | 42.7+/- 8.98 | HDL females<br>mg/dL | 49+/-13 |
| Triglycerides<br>mg/dL | 123.5(83.5-188) | ALT<br>U/L | 28(21.5-39) |
| Gamma-GT<br>U/L | 25(16.5-42.5) | hsCRP<br>mg/mL | 0.56(0.27-1.3) |
| Ferritin Males<br>ng/mL | 167.5(85-234.5) | Ferritin Females<br>ng/mL | 41.5(20-69) |
| Fibrinogen<br>g/L | 306+/-74.7 | Cholesterol<br>mg/dL | 190+/-36.1 |
| Fat mass % | 52+/-7.5 | FFM % | 56.2(41-67) |
| RMR/FFM/Kg | 38.7(33.9-43) | HS at US<br>Grade 1/2/3 (n) | 22/50/8 |
| VAT at US<br>cm | 7.5(6-9.4) | SAT at US<br>cm | 2.6(2.1-3.1) |
| IMTG score | 2.25<br>(2.-3) | MS<br>yes/not (n) | 51/29 |
| IFN-alpha2<br>pg/mL | 120.1+/-24.6 | IFN-gamma<br>pg/mL | 158(57-390) |

The score of IMTG was not different when controlled for gender (Table 2a), while the severity of HS at US was related to the obesity degree (Table 2b). Finally, the score of IMTG was not dependent from the obesity degree (Table 2c).

**Table 2a:** Correlation between IMTG and Gender. Legend of Table 2a: Two-Way cross-tabulation, Pearson chi square, P = 0.6; Total: Number of patients.

| Gender | IMTG scale (scores) |  |  |  | Total |
| --- | --- | --- | --- | --- | --- |
|  | I | II | III | IIII |  |
| Females | 7 | 14 | 16 | 7 | 44 |
| Males | 3 | 10 | 14 | 9 | 36 |

**Table 2b:**
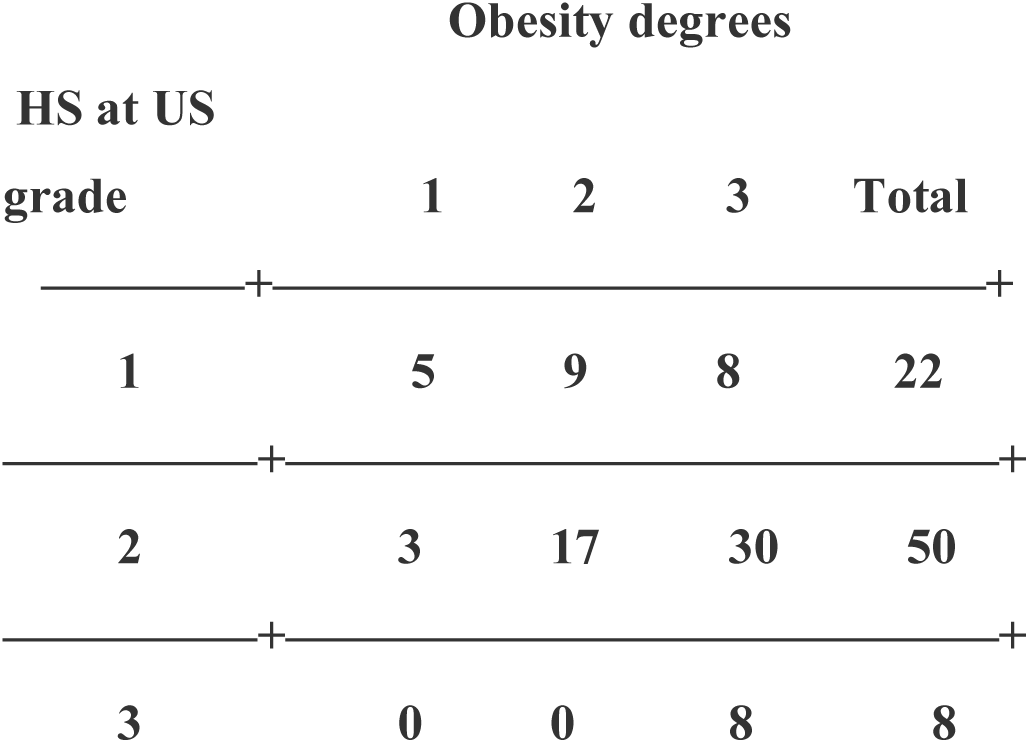
Correlation between Obesity severity and hepatic steatosis grade Obesity degrees. Legend of Table 2b: Two-Way cross-tabulation, Pearson chi2 = 12.5536 P = 0.014; Total: Number of patients.

**Table 2c:**
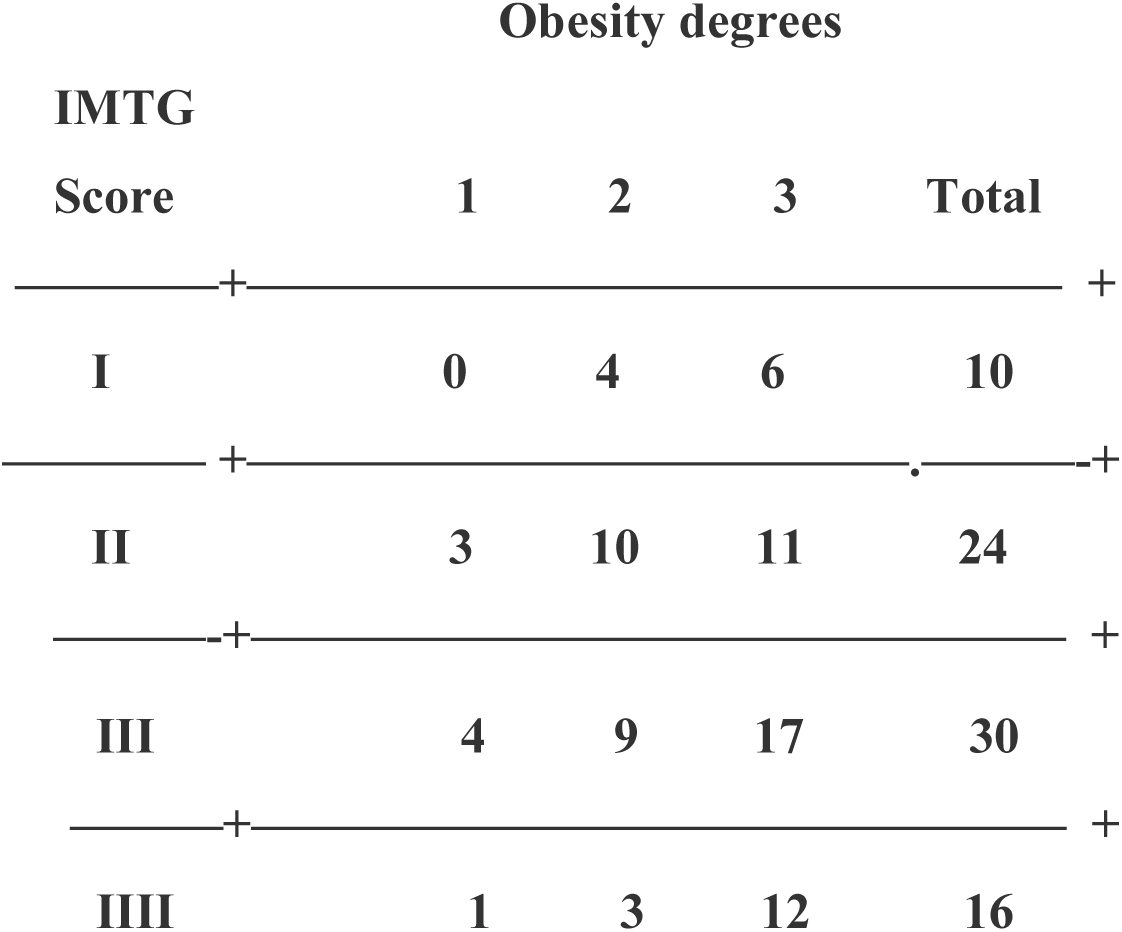
Correlation between Obesity severity and IMTG. Legend of Table 2c: Prevalence of moderate/severe grade of IMTG, i.e. 54 out of 80 patients = 67.5%; Two-Way cross-tabulation, Pearson chi2 = 4.9252, P = 0.553; Total: Number of patients. Hepatic Steatosis (HS) at UltraSonography(US).

### Relationships

First of all, IFN-alpha 2 and IFN-gamma levels were not correlated, p = 0.67, Spearman’ s rank correlation.

IMTG score were negatively predicted by IFN-alpha 2 levels at robust regression and ordered profit regression, Coeff. = −.0151912 Std:Err. = .0043683, t = −3.48, P = 0.001, Conf. Interval = −.0238932−.0064892 and Coeff. = −.0312922, Std.Err. = .0090407, Z = −3.46, P>|z| = 0.001, Conf. Interval. = −.0490116−.0135729, respectively, Figure 1. The residual-versus-fitted plot, Figure 3 of supplemental data, shows that fitted values do not have an obvious trend of failure. Conclusively, there is no problems of heteroskedasticity as residuals appear to have the same variance everywhere. When adjusting for gender and age the prediction of IFN-a2 on IMTG overlapped the previous ones obtained by two methods (robust regression and ordered profit regression), i.e., Coeff. = −.0149346, Std.Err. = .0037878, t = −3.94, P = 0.000, Conf. Interval.=−.0224818-−.0073873.

**Figure 1 Supplem.data.**
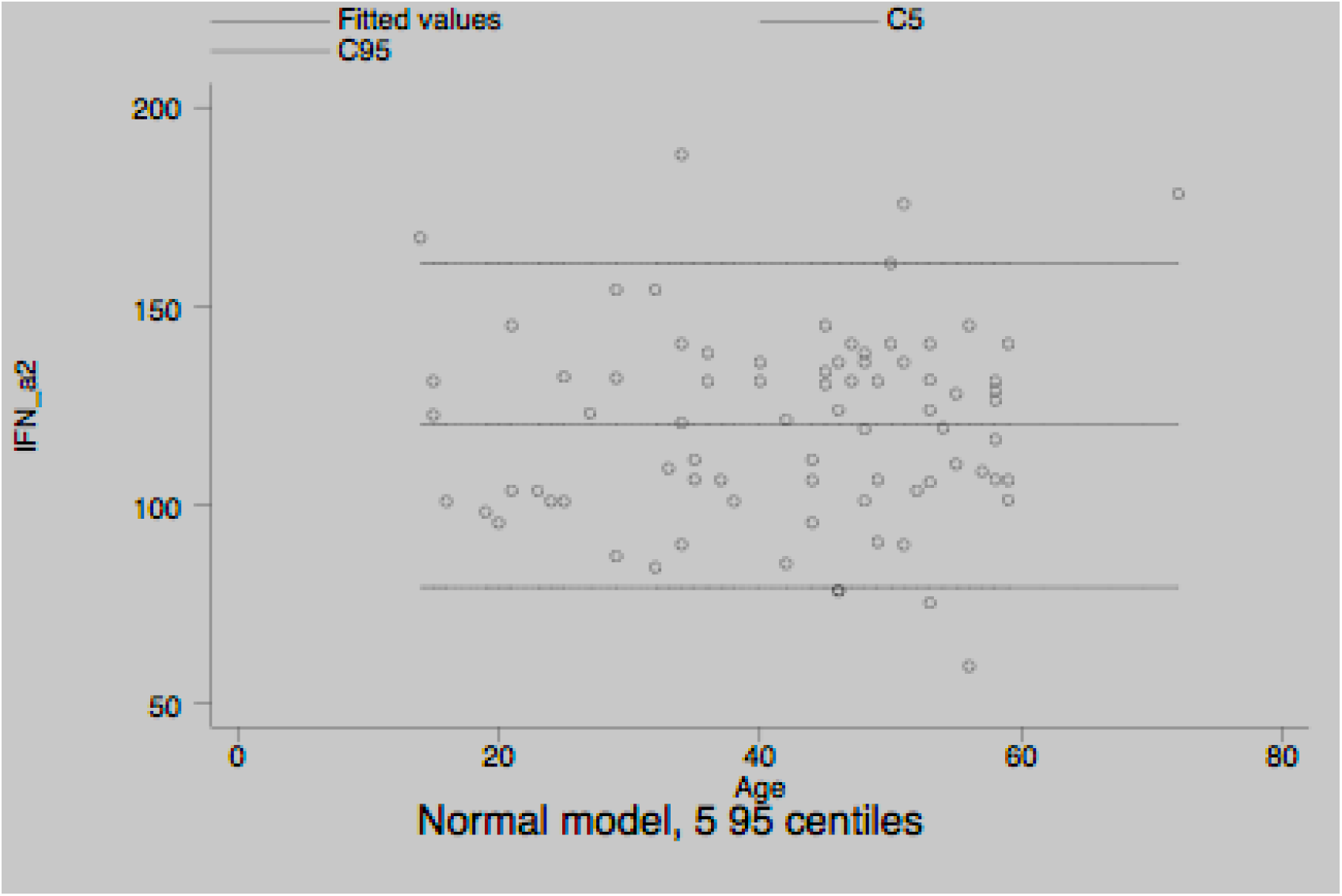
Age-related reference intervals of IFN-a2 in healthy subjects. Legend to Figure 1 Legend to Figure 1 Supplem.data: This is not a “live” graph and therefore cannot be edited. IFN-a2 are expressed in pg/mL

**Figure 2 Supplem.data:**
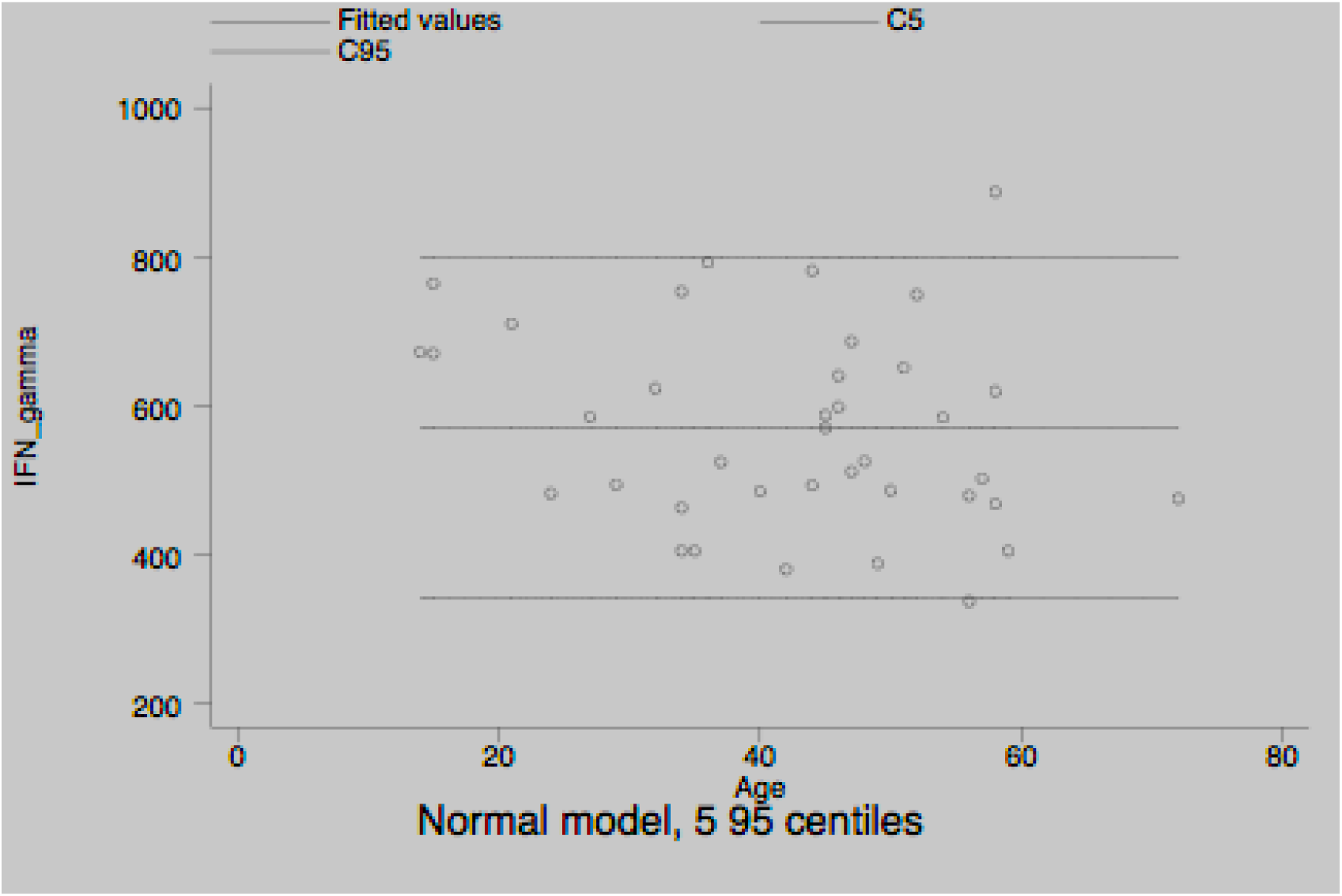
Age-related reference intervals of IFN-gamma in healthy subjects. Legend to Figure 2 Supplem.data: This is not a “live” graph and therefore cannot be edited. IFN-gamma are expressed in pg/mL.

**Figure 3 of Supplem. Data:**
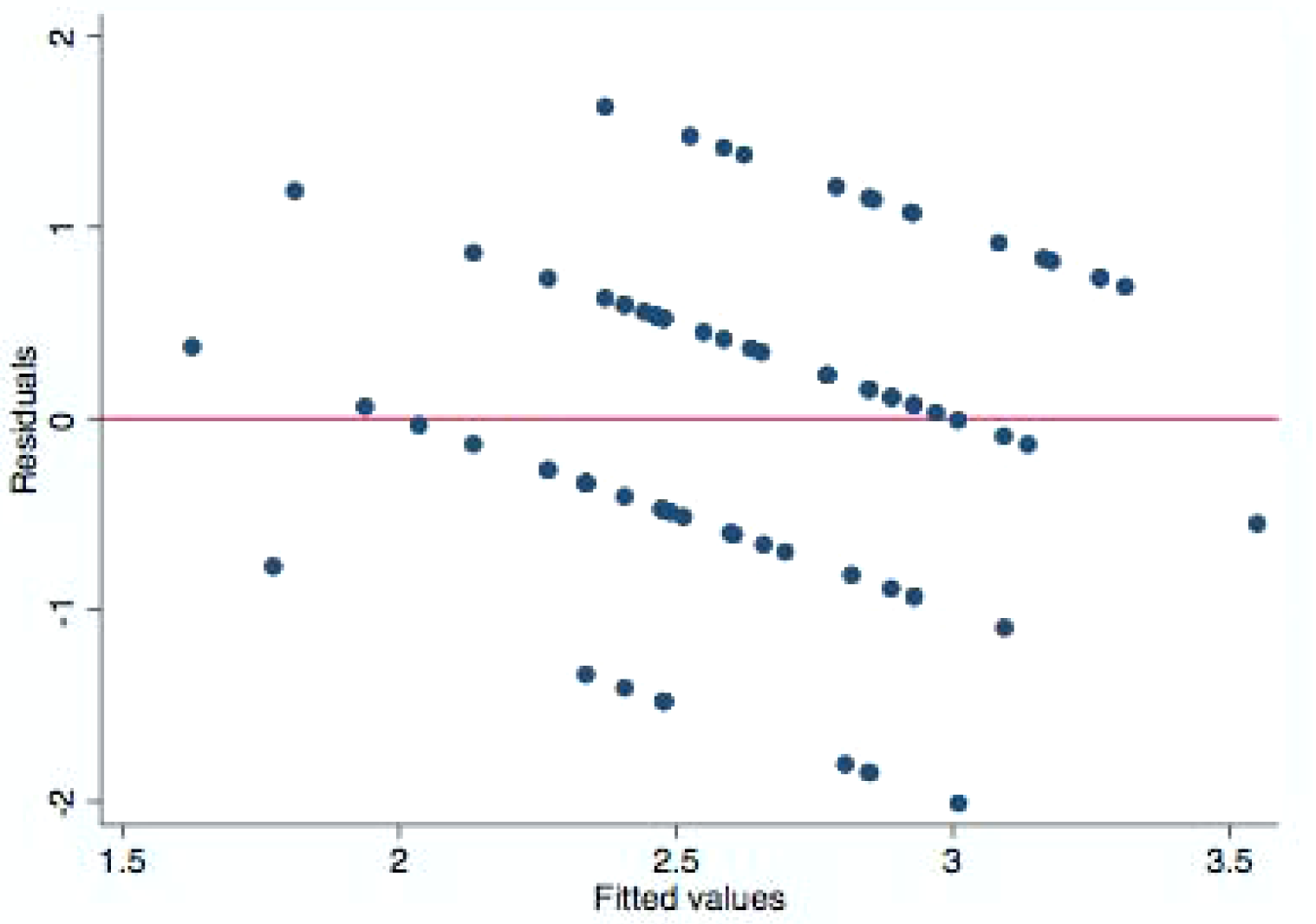
Legend to Figure 3 of Supplem.data: Regression diagnostics, i.e., residuals plotted against the fitted values. This residual plots was used to assess whether the observed error (residuals) was consistent with stochastic error., i.e., differences between the expected and observed values were random and unpredictable. In a well-fitted model, there should be no pattern to the residuals plotted against the fitted values. Ignoring the three outliers at the left side of the graph and one at the right side, we do not see curvature or irregularity in the pattern of the residuals, suggesting a violation of the assumption that IMTG is linear in our independent variable, i.e., IFN-alpha. We might also have seen increasing or decreasing variation in the residuals— heteroskedasticity. Conclusively, any pattern whatsoever indicates a violation of the least-squares assumptions

IMTG grades were not predicted by IFN-gamma levels, analysing them both by LAD regression and order profit regression, i.e., Coeff. = .0000239, Std .Err = .0003695, t = 0.06, P = 0.94995%, Conf. Interval. = −.0007118 -.0007596 and Coeff. = 0000152, Std. Err. = 0004215, z = 0.04, P>|z| = 0.971, Conf. Interval = −.0008109-−.0008412, respectively.

Table 3 shows results of the simultaneous quantile regression (bootstrap method), highlighting that IMTG scores are predicted exclusively by intermediate and upper quantiles of IFN-alpha 2 levels.

**Table 3:** Quantile regression predicting IMTG by IFN-alpha. Legend to Table 3: Simultaneous Quantile regression Bootstrap SE (200 replications). The prediction of IMTG by IFN-alpha2 levels is confined to their intermediate and upper quantiles.

IMTG
|  | Coeff. | Std.Err. | t | P | 95 %CI |
| --- | --- | --- | --- | --- | --- |
| <u>q25</u> |  |  |  |  |  |
| IFN_alpha2 | 0 | .0036305 | 0.00 | 1.000 | -.0072308-.0072308 |
| <u>q50</u> |  |  |  |  |  |
| IFN_alpha2 | -.0174917 | .004081 | -2.58 | 0.08 | -.0302554 -.0047389 |
| <u>q75</u> |  |  |  |  |  |
| IFN_alpha2 | -.0181554 | .0045649 | -3.98 | 0.000 | -.0272471 -.0090637 |

Although IFN-alpha 2 was playing *per se* a significant role in predicting fibrinogen in the mediation method, its role was completely excluded as evident in Table 4.

**Table 4:**
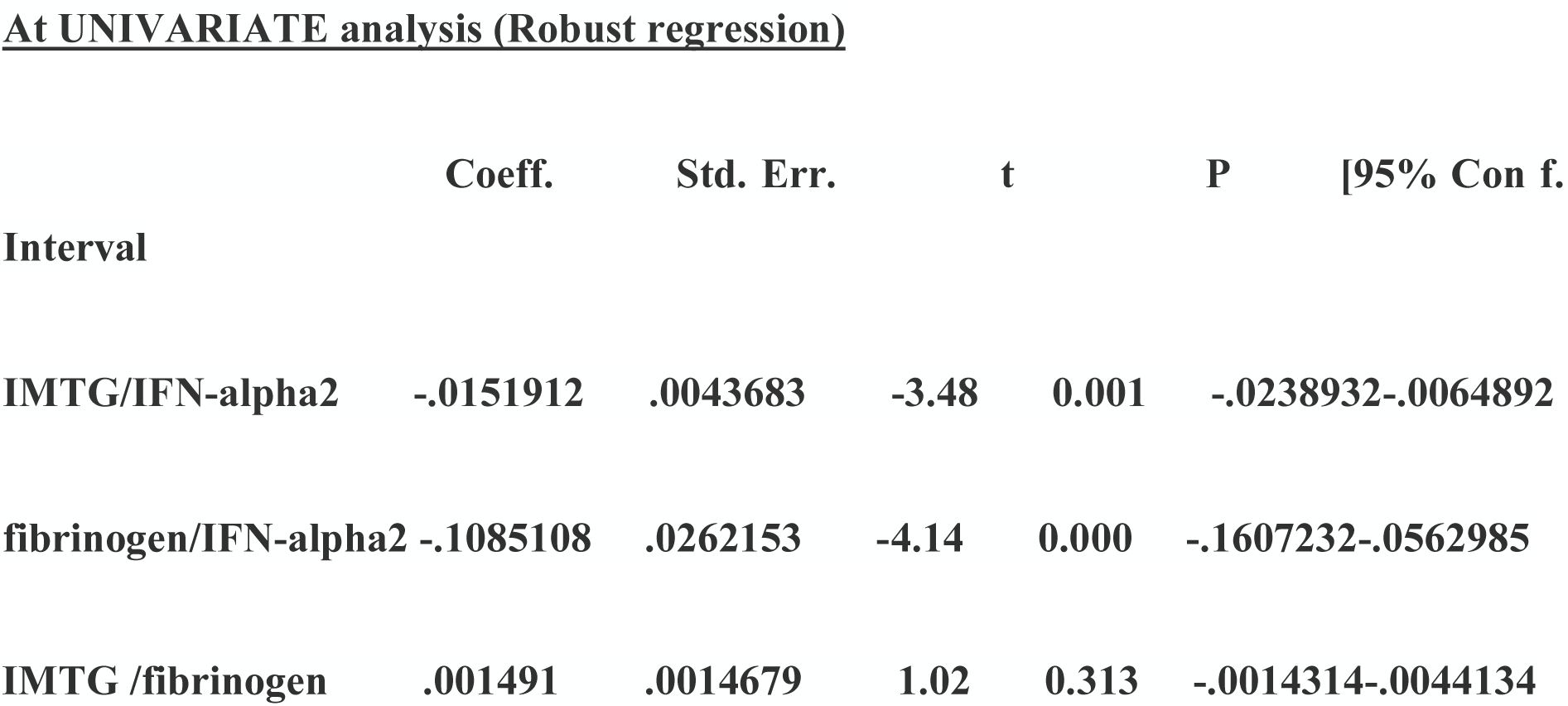
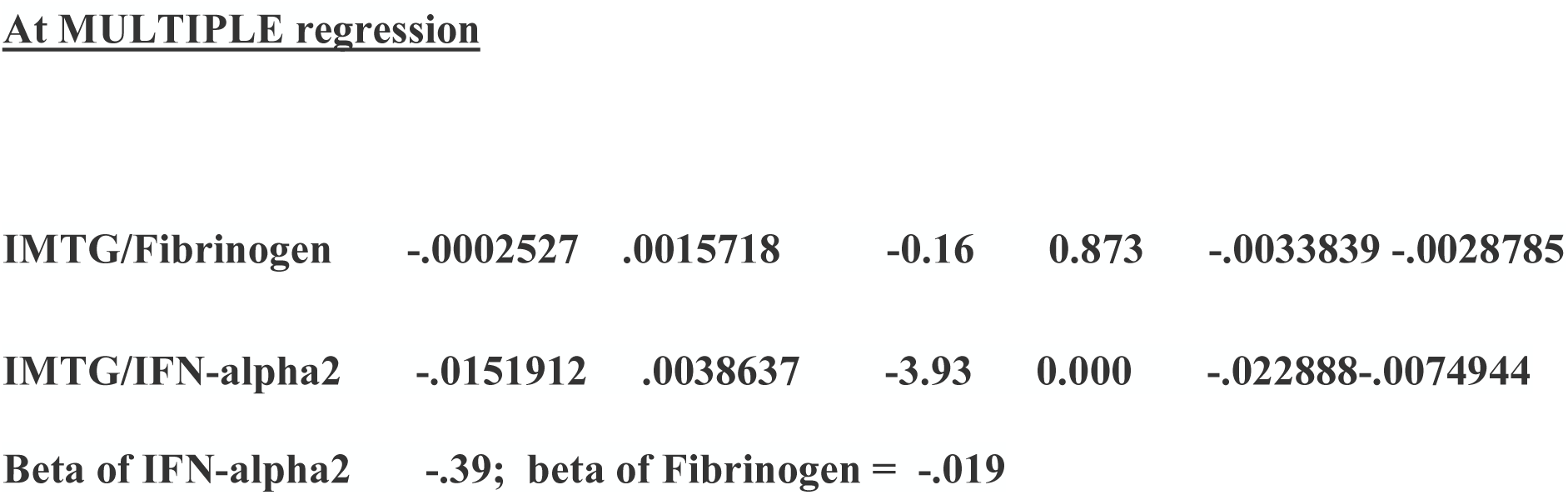
Mediation method. Legend to Table 4: The mediation effect of fibrinogen was excluded (text) The ordered probit regression showed that IMTG was predicted by HS at US as well as HS was predicted by VAT, Table 5, Figure 2.

At UNIVARIATE analysis (Robust regression)
|  | Coeff. | Std. Err. | t | P | [95% Con f. |
| --- | --- | --- | --- | --- | --- |
| Interval |  |  |  |  |  |
| IMTG/IFN-alpha2 | -.0151912 | .0043683 | -3.48 | 0.001 | -.0238932-.0064892 |
| fibrinogen/IFN-alpha2 | -.1085108 | .0262153 | -4.14 | 0.000 | -.1607232-.0562985 |
| IMTG /fibrinogen | .001491 | .0014679 | 1.02 | 0.313 | -.0014314-.0044134 |
| **At MULTIPLE regression** |  |  |  |  |  |
| IMTG/Fibrinogen | -.0002527 | .0015718 | -0.16 | 0.873 | -.0033839 -.0028785 |
| IMTG/IFN-alpha2 | -.0151912 | .0038637 | -3.93 | 0.000 | -.022888-.0074944 |

Beta of IFN-alpha2 -.39; beta of Fibrinogen = -.019

The ordered probit regression showed that IMTG was predicted by HS at US as well as HS was predicted by VAT, Table 5 and Figure 2.

**Table 5:**
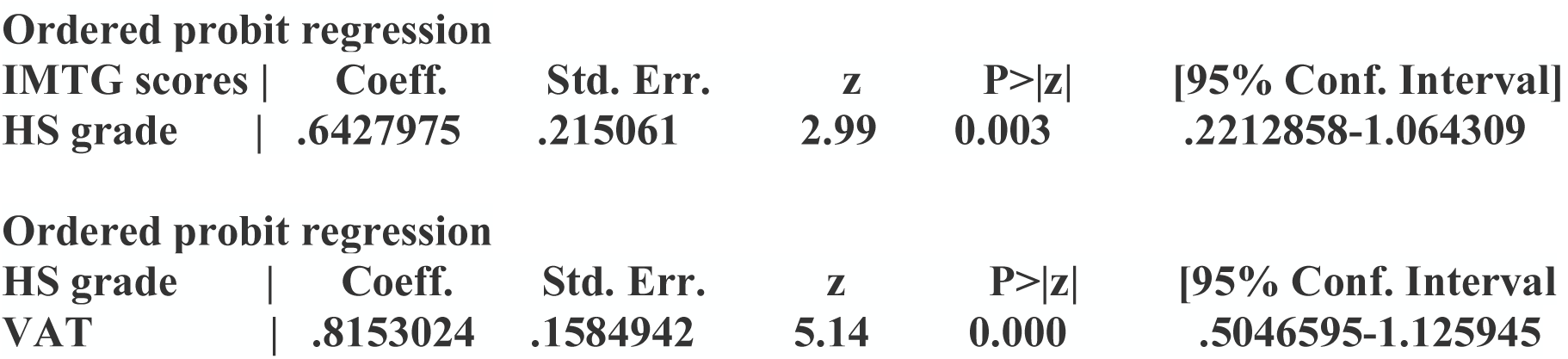
Legend to table 5: In these two regressions there is a suspicion of a confounding variable or covariate that is VAT.

|  |  |  |  |  |  |
| --- | --- | --- | --- | --- | --- |
| IMTG scores \| | Coeff. | Std. Err. | z | P>\|z\| | [95% Conf. Interval] |
| HS grade \| | .6427975 | .215061 | 2.99 | 0.003 | .2212858-1.064309 |
| **Ordered probit regression** |  |  |  |  |  |
| HS grade \| | Coeff. | Std. Err. | z | P>\|z\| | [95% Conf. Interval] |
| VAT \| | .8153024 | .1584942 | 5.14 | 0.000 | .5046595-1.125945 |

To clarify this important aspect, instrumental-varables regression excluded the role of the confounding variable (VAT) as mediator between IMTG and HS, as evident in Table 6. At multivariate analysis, among VAT, SAT, WC, WHR, BMI only VAT predicted IMTG, i.e., Coeff. = .1384446, Std.Err. = .0582611, t = 2.38; P>|t| = 0.020, 95% Conf. Interval].0223568-.2545323.

**Table 6:**
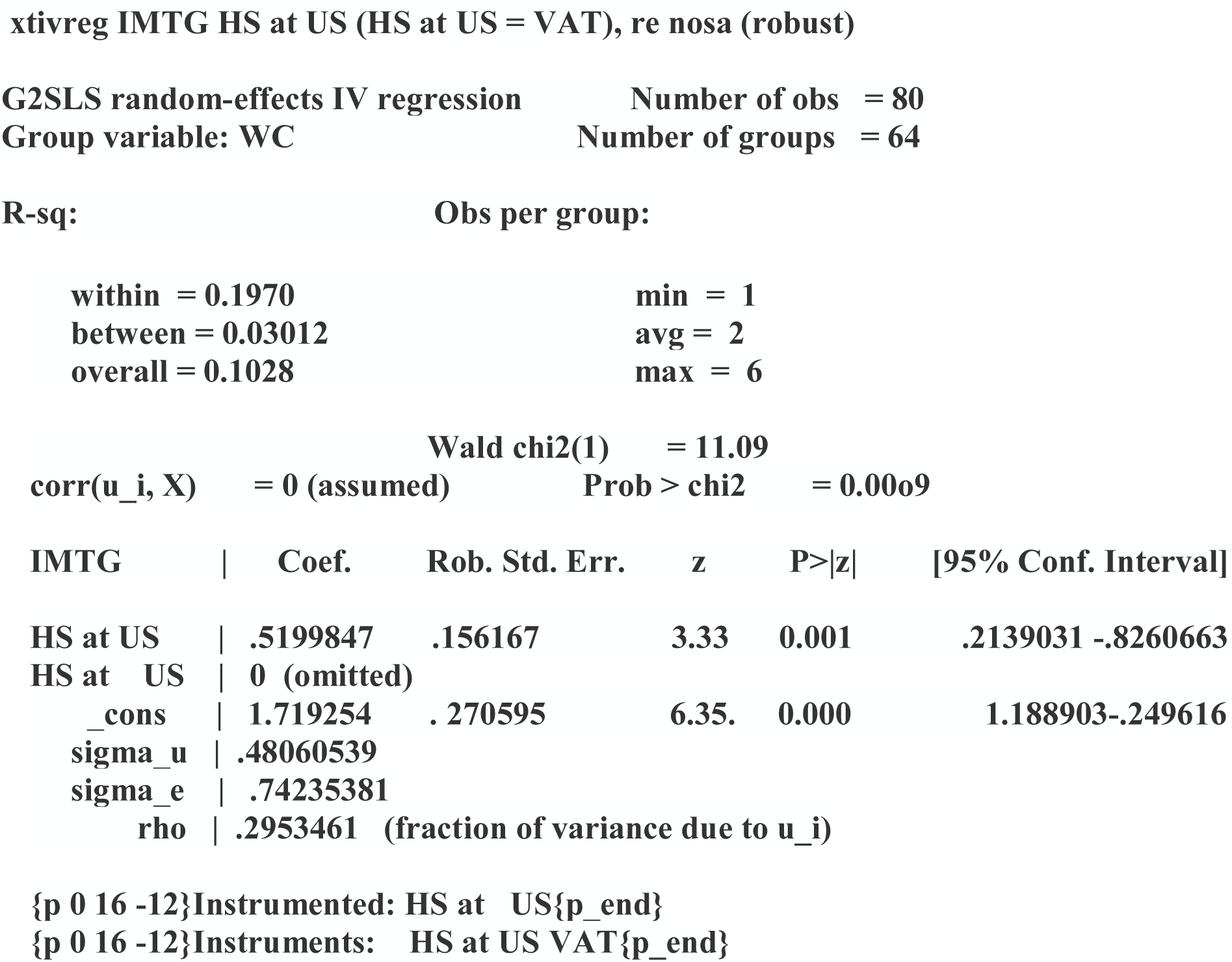
Instrumental-variables regression for panel data (Baltagi-Chang estimator) xtivreg IMTG HS at US (HS at US = VAT), re nosa (robust) Legend to Table 6. The instrument (VAT) cannot be correlated with the dependent (IMTG) in the explanatory equation. In other words, the instrument cannot suffer from the same problem as the original predicting variable (IMTG predicted by HS at US). If this condition is met, then the instrument is said to satisfy the exclusion restriction. As grouping variable was chosen an index of visceral adiposity, i.e., WC. IMTG, intramusclolar triglycerides; HS at US, hepatic steatosis at Ultrasonography; VAT, visceral adipose tissue.

|  |  |
| --- | --- |
| **G2SLS random-effects IV regression** | **Number of obs = 80** |
| **Group variable: WC** | **Number of groups = 64** |
| **R-sq:** | **Obs per group:** |
| within = 0.1970 | min = 1 |
| between = 0.03012 | avg = 2 |
| overall = 0.1028 | max = 6 |

|  |  |  |
| --- | --- | --- |
|  | Wald chi2(1) | = 11.09 |
| corr(u_i, X) | = 0 (assumed) | Prob > chi2 = 0.0009 |

| IMTG | Coef. | Rob. Std. Err. | z | P>\|z\| | [95% Conf. Interval] |
| --- | --- | --- | --- | --- | --- |
| HS at US | .5199847 | .156167 | 3.33 | 0.001 | .2139031 -.8260663 |
| HS at US | 0 (omitted) |  |  |  |  |
| _cons | 1.719254 | .270595 | 6.35 | 0.000 | 1.188903-.249616 |
| sigma_u | .48060539 |  |  |  |  |
| sigma_e | .74235381 |  |  |  |  |
| rho | .2953461 | (fraction of variance due to u_i) |  |  |  |

{p 0 16 -12}Instrumented: HS at US{p\_end} {p 0 16 -12}Instruments: HS at US VAT{p\_end}

No prediction of IMTG by HOMA, HOMA%B and QUICKI (P = 0.56, 0.15 and 0.71 respectively). The finding concerning no link between IR and IMTG was confirmed by a more powerful tool evaluating whether IMTG might be predicted by HOMA, i,e, Coef. = .0219949; Std.Err. = .0367399 z = 0.60; P>|z| >0.549; Conf. Interval = -.050014− .0940038; ordered profit regression, robust method.

There was no prediction of IMTG by fat mass, FFM and RMR/FFM ratio (P = 0.550, 0.232 and 0.069, respectively, evaluated as ordered profit regression). By the same technique, HS at US was not predicted by IFN-alpha levels (P = 0.079). As expected, at univariate analysis HOMA%B was predicted by HOMA: Coef. 8.200134, Std.Err. = 1.072445, t= 7.65, P>|t| = 0.000, 95% Conf. Interval.= 6.06506-10.33521. Studying the hidden relationships between various variables comprehending anthropometric measures, fat and free-fat masses, US features of central, peripheral fat and intramuscular distribution, it is confirmed the link between IFN-a2 and IMTG (as evident in factor 3) as well as other parameters, Table 7.

**Table 7:**
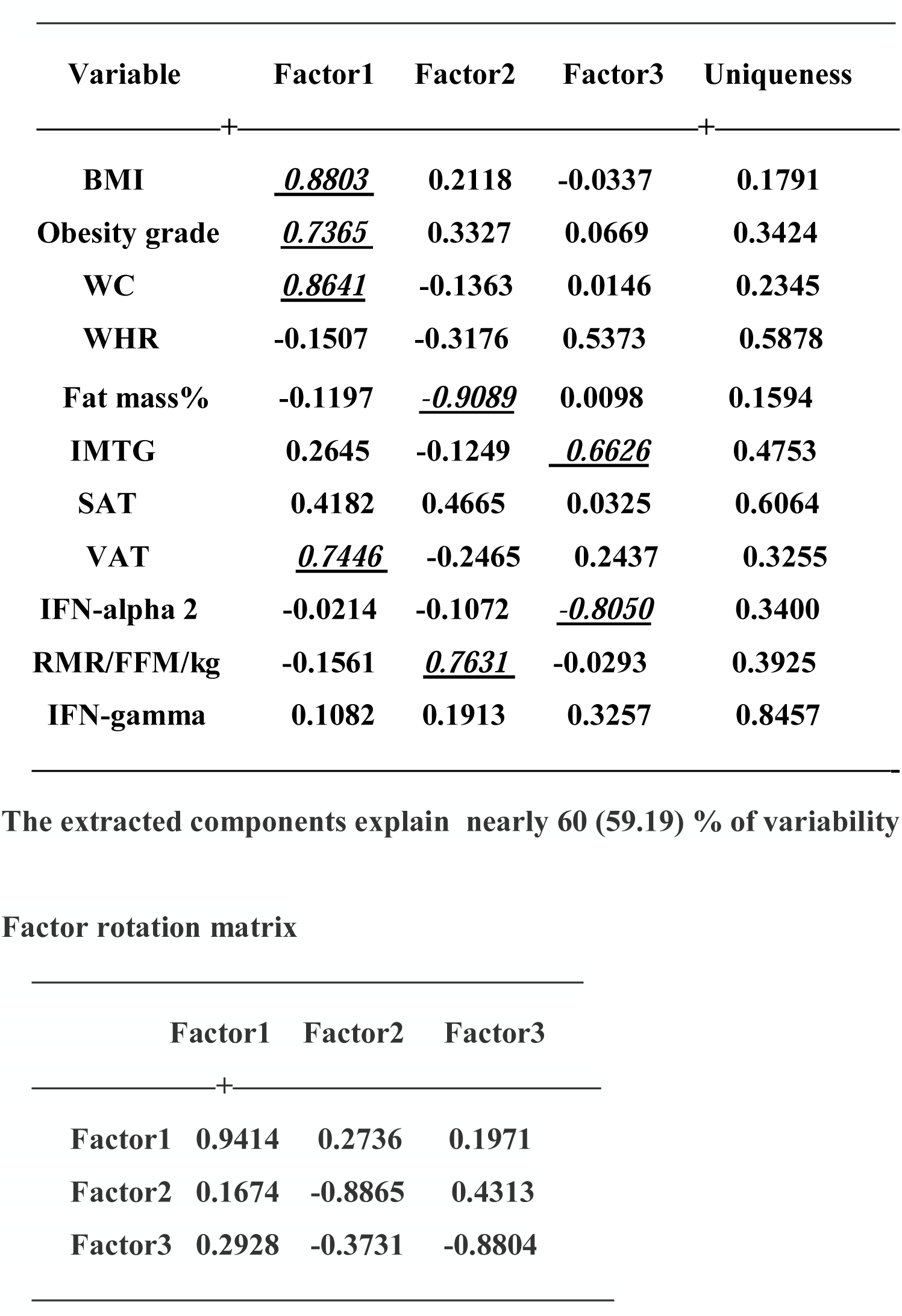
Factor analysis Rotated factor loadings (pattern matrix) and unique variances. Legend to Table 7: The critical value was calculated by doubling Pearson’s correlation coefficient for 1% level of significance (5.152)/square root of patients minus 2 (78), i.e., 0.583. In underlined it will be shown the main components for any single factor, with a value superior to the critical one. HS was excluded due to collinearity with IMTG and VAT. The link between IFN-a2 and IMTG (factor 3) as well as significative parameters in factors one and two are shown in italics and underlined text.

| Variable | Factor1 | Factor2 | Factor3 | Uniqueness |
| --- | --- | --- | --- | --- |
| BMI | <u>0.8803</u> | 0.2118 | -0.0337 | 0.1791 |
| Obesity grade | <u>0.7365</u> | 0.3327 | 0.0669 | 0.3424 |
| WC | <u>0.8641</u> | -0.1363 | 0.0146 | 0.2345 |
| WHR | -0.1507 | -0.3176 | 0.5373 | 0.5878 |
| Fat mass% | -0.1197 | <u>-0.9089</u> | 0.0098 | 0.1594 |
| IMTG | 0.2645 | -0.1249 | <u>0.6626</u> | 0.4753 |
| SAT | 0.4182 | 0.4665 | 0.0325 | 0.6064 |
| VAT | <u>0.7446</u> | -0.2465 | 0.2437 | 0.3255 |
| IFN-alpha 2 | -0.0214 | -0.1072 | <u>-0.8050</u> | 0.3400 |
| RMR/FFM/kg | -0.1561 | <u>0.7631</u> | -0.0293 | 0.3925 |
| IFN-gamma | 0.1082 | 0.1913 | 0.3257 | 0.8457 |

Factor rotation matrix
|  | Factor1 | Factor2 | Factor3 |
| --- | --- | --- | --- |
| Factor1 | 0.9414 | 0.2736 | 0.1971 |
| Factor2 | 0.1674 | -0.8865 | 0.4313 |
| Factor3 | 0.2928 | -0.3731 | -0.8804 |

## DISCUSSION

This study was designed to investigate any correlation of IFN-alpha 2a and IFN-gamma to IMTG in obese patients with US-detected HS or NAFLD. Stating the major findings of our study, we should drawn attention on: j) serum concentrations of IFN-alpha2 were increased, while serum levels of IFN-gamma were decreased confronted with those of controls; jj) the severity of IMTG, revealed at US as Heckmatt scores (I-IV) was negatively predicted by IFN-alpha2 serum concentrations; jjj) IMTG scores were not predicted by serum levels of IFN-gamma; jjjj) IMTG scores were predicted by HS severity ascertained by a US scale (grades 1-3); jjjjj) HS severity was predicted by VAT but the latter was not instrumental to IMTG.

To try to explain the possible mechanisms of the core finding, i.e., the inverse association between IFN-alpha2 and IMTG we can focus on serum lipid profile creating sort of parallelism between other situations/diseases in which increased concentrations of this cytokine were found and our obese patients.

Previous experiments in vitro showed that IFN-alpha inhibits lipoprotein lipase (LPL) activity directly in patients with condylomata acuminata treated with interferon alpha-n1 or indirectly by inducing other monokines such as Interleukin-1 and tumor necrosis factor alpha (31, 32). What is more, strong evidence has indicated that apo C III suppresses LPL-induced hydrolysis of VLDL-triglyceride ending up in hypertriglyceridemia and an increase in apo E levels (33.) Generally, increase in the apo C-III and apo levels mirrors decrease in VLDL clearance. In addition, increase in lipogenesis and VLDL secretion in the liver by IFN-alpha may contribute to hypertriglyceridemic subjects, since IFN-alpha stimulates lipogenesis in some cultured hepatocytes (34-36) and in patients suffering from HCV-related chronic hepatitis on IFN-alpha therapy (37). Coming back to the role of IMTG, recent reports indicate that the elevated IMTG content found in obese women was not due to an up-regulation of key lipogenic proteins or to the suppression of lipolytic proteins. At this point perils-in could play an essential role. In fact, phosphorylation of qperilipin is essential for the mobilization of fats in adipose tissue. Unfortunately, the impact of a low perilipin protein abundance relative to the amount of IMTG in obesity remains to be clarified (38).

Although IMTG synthesis rates were previously related to insulin sensitivity (2), we authors did not find a link between IMTG and IR, but it is noteworthy to stress that we did not evaluate IR by glucose clamp technique but surrogate markers, even though quite reliable (39).

Relating our findings to those of similar studies, we emphasise that in a cross-sectional study on 42 HIV-1-positive men on antiretroviral therapy, 27 of whom had symptoms of lipodystrophy (LD), defined by computed tomography scan, serum IFN-alpha was markedly increased in LD-positive compared with LD-negative men and controls. A significant positive correlation was found between accumulation of IFN-alpha and increased levels of cholesterol, TG, VLDL cholesterol, VLDL TG, ApoB and ApoB-ApoA1 ratio (40). In the opposite sense was the study of Teran-Cabanillas *et al*, who found that obese subjects showed a decreased ability to produce IFN-alpha but also IFN-βeta in response to TLR ligands; this response was associated with increased basal levels of SOCS3 but not SOCS1(41)

Concerning the lower levels of IFN-gamma in our population confronted with those of controls, we recognise that the lack of physical evaluation does not permit a comparison with results by Schmidt *et al* (42), who found that general obesity and central obesity are associated with significantly elevated IFN-gamma-levels, but in obese subjects physical activity may lower levels and thus reduce pro-inflammatory effects of cytokines that may link obesity, insulin resistance and diabetes. Vice versa, looking at the levels of IFNs in our obese and control arms, they overlap with those expressed into literature (43).

The association between IMTG scores and HS at US grades could hypothetically explained on the ground of common mechanisms leading to ectopic fat storage and utilisation. Indeed, the reason for which there is a lack of link between IMTG and VAT remains a point to be further clarified: this no association is in agreement with the absence of correlation between IMTG scores and IR and in contrast with the apparent but not confirmed, when adjusted for VAT presence, association between IMTG and HS at US. In other words, the link between IMTG and HS is not mediated by the entity of visceral adiposity, evaluated as VAT. This finding bears a certain novelty. Further study is needed to clarify this aspect, at the light that NAFLD is considered a CV risk factor (44) and could lead to hepatocarcinoma.

The importance of relation between IFNs and VAT is underlined by a recent work stressing that type 1 IFN signature gene expression in VAT correlates with both adipose tissue and systemic IR in obese individuals, which is represented by ADIPO-IR and HOMA2-IR, respectively, and defines two subgroups with different susceptibility to IR (45). Finally, up-to-date results in mice provide genetic evidence that plasmacytoid dendritic cells via type I IFNs, regulate energy metabolism and promote the development of obesity (46). An possible mechanism explaining the latter event could be the signaling pathway activated by type I IFNs.

This consists of a series of events: Phosphorylation and activation of two enzymes of the JAK family, TYK2 which is associated with IFNAR1 and JAK1 associated to IFNAR2; phosphorylation by the activated JAK kinases of key transcription factors, namely STAT1 and STAT2, (47). Dysregulation of this pathway leads to increased angiogenesis which modulates adipogenesis, expanding fat depots during weight gain.

Although fatty acids induce type I IFN responses in murine hepatocytes/macrophages and exposure to a high-fat diet elicited type I IFN-regulated gene expression in the liver of wild-type mice, modulating susceptibility to metabolic or hepatic disease (48), we were not able to confirm this important link of IFN-alpha2 with HS in our population.

Among limitations to study, we firstly acknowledge that our was an observational study in which a clear relation of cause and effect is not possible to find; secondly the hypothetical mechanisms are far to be elucidated, being mechanistic studies not carried out; thirdly, although IMTG was associated with metabolic risk factors, most of these associations were reported to be lost after adjustment for BMI or VA, even though a unique association was found to remain for metabolic syndrome in women and lipids in men (49); still the APT III criteria (50) for assessing metabolic syndrome is not completely accepted due to differences in cut-off points; finally, the sample size was not that large.

### Conclusion

The complex interplay between cytokines and ectopic fat excess is hopefully enriched by the observation that in obese patients with NAFLD the serum levels of IFN-alpha2 are inversely related to IMTG scores, differently from IFN-gamma levels that are not associated with severity of this ectopic storage. More controlled research is needed to confirm these preliminary data.

